# Transparent and Reproducible Research Practices in the Surgical Literature

**DOI:** 10.1101/779702

**Authors:** Taylor Hughes, Andrew Niemann, Daniel Tritz, Kryston Boyer, Hal Robbins, Matt Vassar

**Affiliations:** Oklahoma State University Center for Health Science; Kansas City University of Medicine and Biosciences; Oklahoma State University Center for Health Sciences; Oklahoma State University Medical Center

## Abstract

Previous studies have established a baseline of minimal reproducibility in the social science and biomedical literature. Clinical research is especially deficient in factors of reproducibility. Surgical journals contain fewer clinical trials than non-surgical ones, suggesting that it should be easier to reproduce the outcomes of surgical literature. In this study, we evaluated a broad range of indicators related to transparency and reproducibility in a random sample of 300 articles published in surgery-related journals between 2014 and 2018. A minority of our sample made available their materials (2/186, 95% C.I. 0–2.2%), protocols (1/196, 0–1.3%), data (19/196, 6.3–13%), or analysis scripts (0/196, 0–1.9%). Only one study was adequately pre-registered. No studies were explicit replications of previous literature. Most studies (162/292 50–61%) declined to provide a funding statement, and few declared conflicts of interest (22/292, 4.8–11%). Most have not been cited by systematic reviews (183/216, 81–89%) or meta-analyses (188/216, 83–91%), and most were behind a paywall (187/292, 58–70%). The transparency of surgical literature could improve with adherence to baseline standards of reproducibility.

## Introduction

Reproducibility is the “ability of a researcher to duplicate the results of a prior study using the same materials as were used by the original investigator.”^1^ Reproducibility is crucial to scientific advancement. Estimates suggest that less than 50% of clinical research studies are reproducible.^2–5^ This widespread and pervasive issue has been dubbed a reproducibility crisis by scientists.^6^ In cancer biology, researchers attempted to reproduce the findings of 50 high-impact publications but had to abandon 32 of these planned reproductions owing to a lack of sufficiently detailed protocols and materials.^7^ Of the 18 remaining attempts, 10 have been completed, of which 5 were largely reproducible, 3 were inconclusive, and 2 were negative.^8^ In psychology, a large-scale reproducibility investigation was conducted on 100 experimental and correlational studies published in prominent psychology journals.^9^ The mean effect size of the reproduced studies was half the magnitude of the original studies. Further, whereas 97% of the original study results were statistically significant, only 36% of results from the reproduced studies reached statistical significance. Investigators of this study found that 47% of the original effect sizes fell within the 95% confidence interval of the reproduced study’s effect size. These findings suggest a reproducibility deficit across disparate fields of study. This problem becomes especially important in clinical research, where participants enroll in clinical trials and expose themselves to therapies with potential risks to improve treatment options for others.

In surgery research, initiatives to promote reproducibility are ongoing. As examples, the *International Journal of Surgical Protocols* publishes research protocols for all surgical specialties. Reporting standards, such as Strengthening the Reporting of Cohort Studies in Surgery (STROCSS)^10^ and Preferred Reporting of Case Series in Surgery (PROCESS),^11^ have been developed to improve the completeness of reporting of surgical studies. The IDEAL Collaboration seeks to improve the quality of surgical research by including recommendations related to reproducibility.^12^ Editorials also have been published in academic journals to alert surgeons to the reproducibility problem. One article published in the *Journal of Thoracic and Cardiovascular Surgery* describes the likely effects of the new National Institutes of Health’s rigor and reproducibility requirements on surgery research. In this editorial, Lawson notes, “we as surgeons are particularly qualified to identify areas in need of improvement in the care for our patients, and we must continue to perform clinical and basic research to move our field into the future.”^13^ A second editorial^14^ published in *Plastic and Reconstructive Surgery* calls for improved reproducibility of clinical studies, given the proliferation of big data and, by virtue, increased production of clinical database studies. Aside from these initiatives, very little is known about the state of reproducibility and transparency in surgery research.

## Methods

This is a cross-sectional observational study design with data extraction performed in a blind and duplicate fashion. We employed similar methodology to Hardwicke et al.,^15^ with minor modifications. We reported each study in accordance with guidelines for meta-epidemiological methodology research.^16^ This research was observational and did not include any human subjects or study participants, making it exempt from institutional review board approval prior to initiation.^17^ The materials, protocol, and raw data used for this study will be uploaded to the Open Science Framework for public access (https://osf.io/n4yh5/).

### Journal and Study Selection

We used the U.S. National Library of Medicine catalog to search for all journals using the subject terms tag Surgery[ST]. This search was performed on 05/29/2019. The inclusion criteria required that journals were in “English” and “MEDLINE indexed.” The list of journals in the U.S. National Library of Medicine catalog then were extracted by the electronic or linking International Standard Serial Number (https://osf.io/t83cy/), which was used to search PubMed for all publications by these journals. The list of publications included those published between January 1, 2014, and December 31, 2018. From this list, 300 publications were randomly sampled to extract their data (https://osf.io/r8tx2/). We randomized all publications within the list to ensure availability of additional publications during the data extraction process, but they were not needed.

### Data Extraction

A pilot-tested Google Form was created based on the one provided by Hardwicke et al.,^15^ with additions. This form prompted coders to identify whether a study had the necessary information to be reproducible (Supplement 1). The data extracted varied based on the study design. Studies with no empirical data (e.g., editorials, commentaries [without reanalysis], simulations, news, reviews, and poems) were excluded. On our form, we changed the original item for identifying the impact factor of a specific year to two new items, one identifying the 5-year impact factor and one identifying the most recent impact factor. We also expanded the options for study design to include cohort, case series, secondary analysis, chart review, and cross-sectional designs. Finally, we expanded the funding options to be more specific to university, hospital, public, private, industry, or non-profit.

Two authors extracted data from the 300 articles in duplicate and blinded fashion. Prior to data extraction, they underwent a full day of training to ensure reliability between authors. The training began with an in-person session to review the study design, protocol, Google Form, and location of the information in two articles. The authors were given three example articles from which to extract data. Following extraction, the pair reconciled any differences between them. This training session was recorded and posted online for reference. Prior to extracting data from all studies, the two authors extracted data from the first 10 articles in their respective list, followed by a final consensus meeting. Data extraction on the remaining 290 articles was then conducted. A final consensus meeting was held by the pair to resolve disagreements. A third author was available for adjudication, if necessary.

### Analysis Plan

We report descriptive statistics for each category using Microsoft Excel. For each analyzed characteristic, 95 perfect confidence intervals were generated.

## Results

### Sample Characteristics

Our search of the U.S. National Library of Medicine database identified 409 surgery journals, with 147 fitting the inclusion criteria. A PubMed search of those 147 journals identified 681,637 publications, with 154,441 published within the specified time-frame. From this list, we randomly sampled 300 publications. Of the 300-article sample, eight (2.7%) full-text documents were unable to be accessed. The remaining coded articles came from a broad range of journals, with a median 5-year impact factor of 2.727 and a most recent impact factor median of 2.471 (data from 2017 and 2018 when available). Impact factor data was unavailable for 26 coded articles. Table 1 lists more sample characteristics.

**Table 1:**
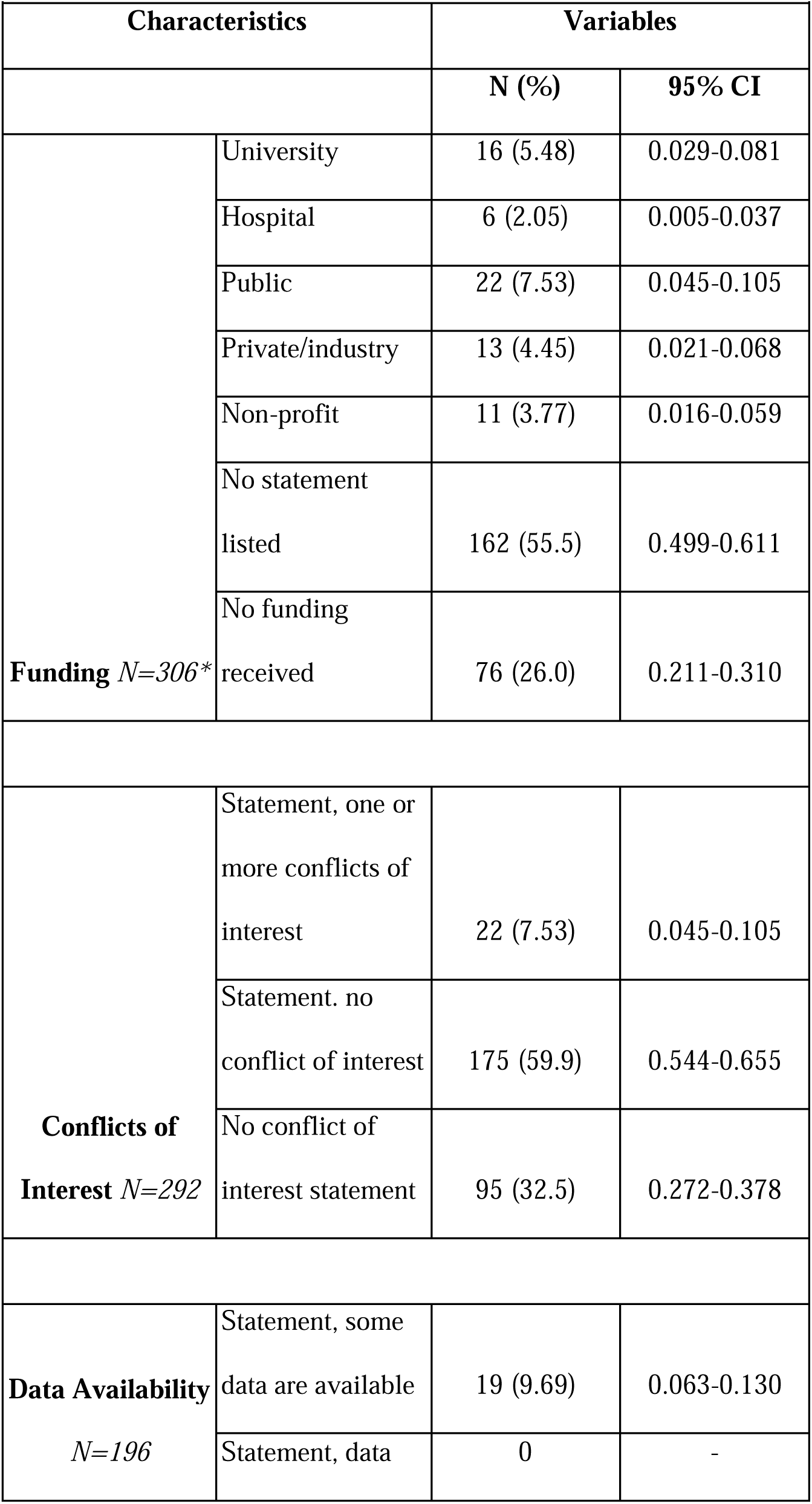
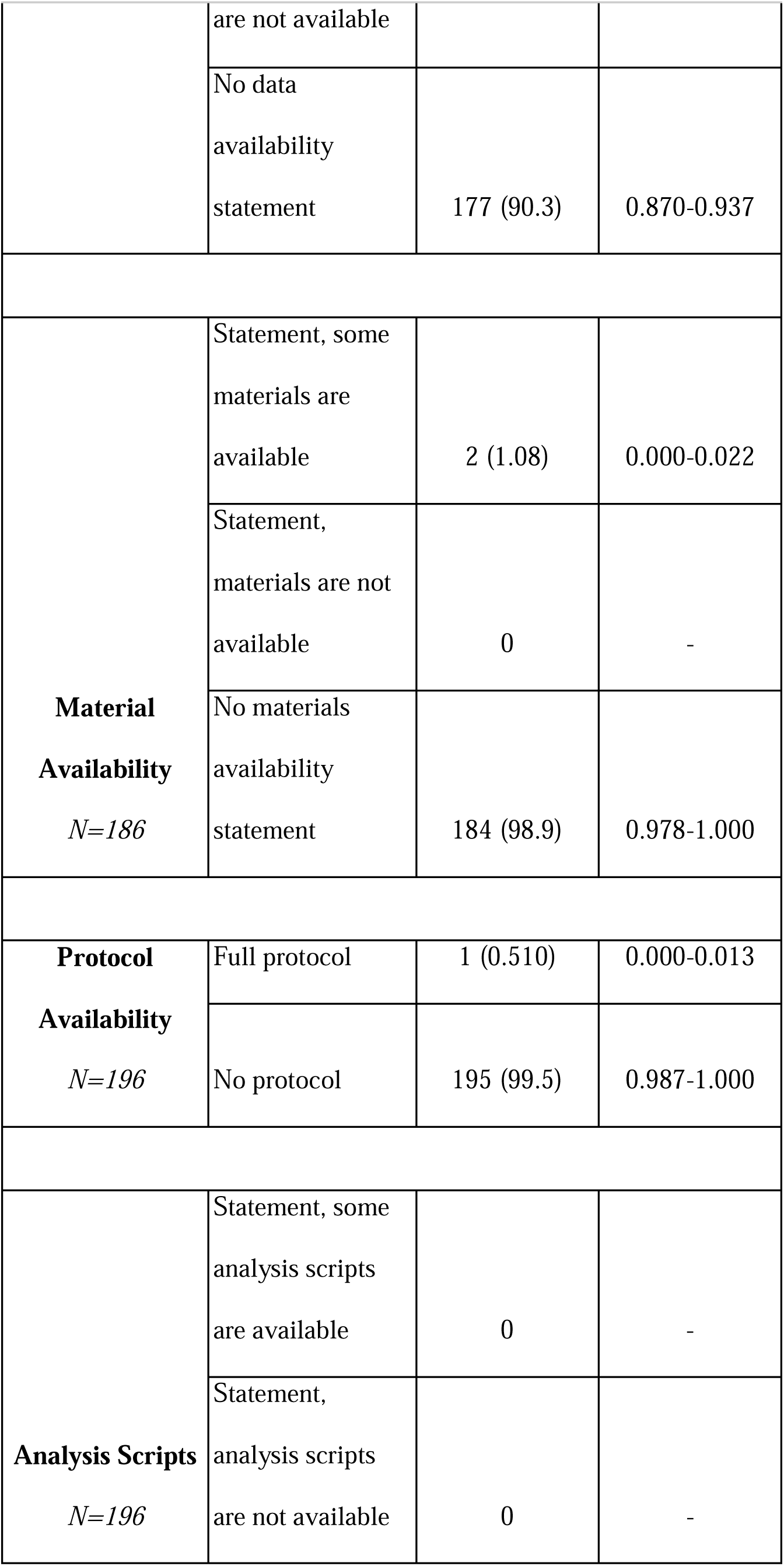
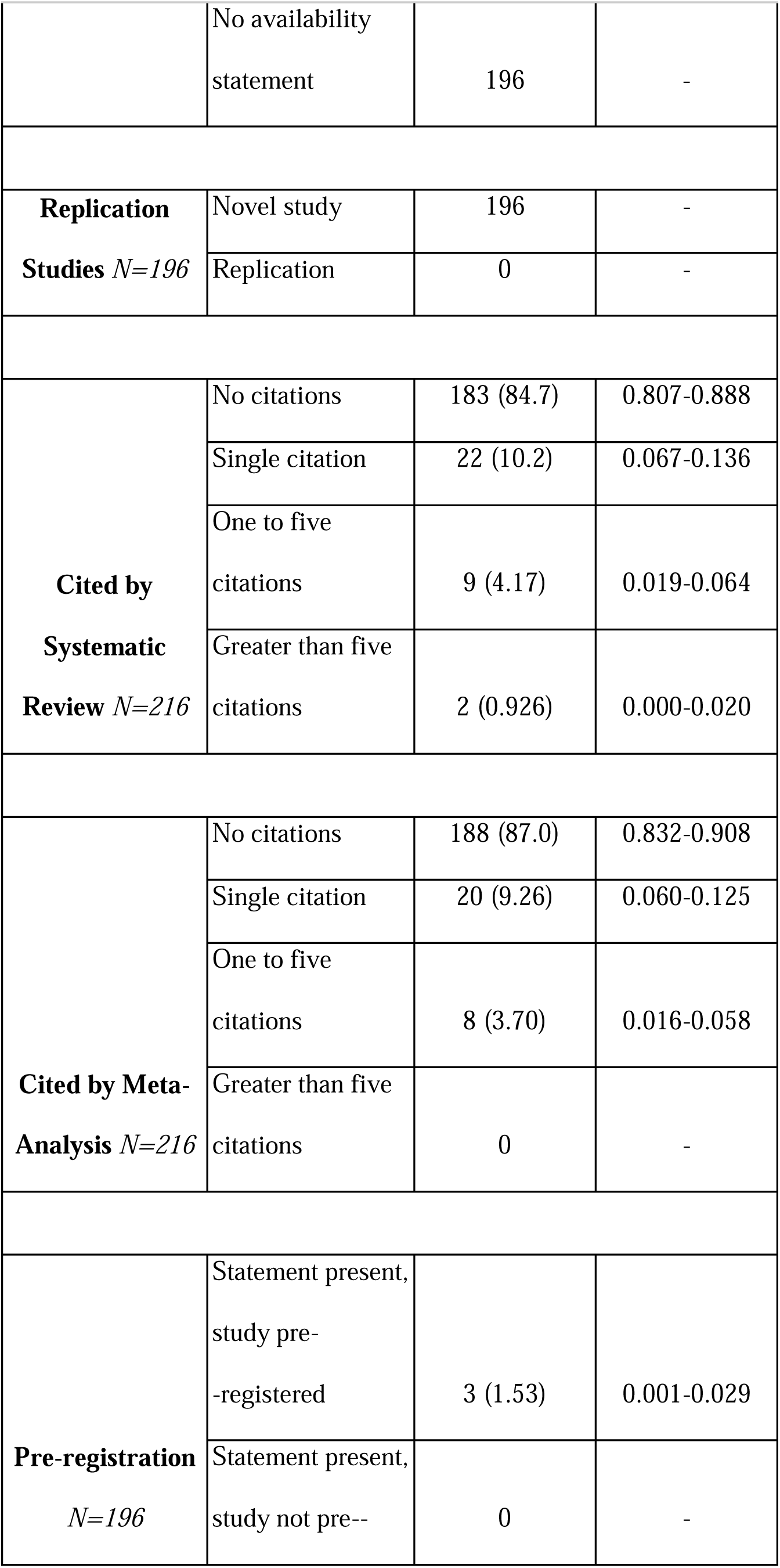
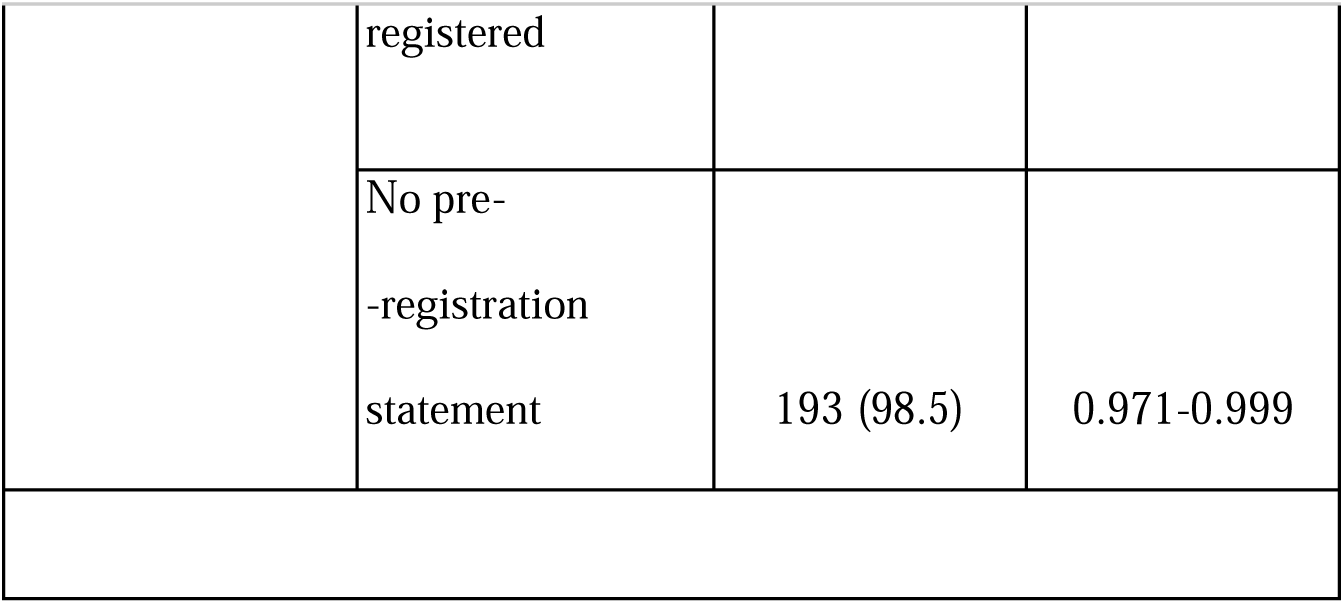
Characteristics of Reproducibility in Surgery Studies.

### Access to Articles

Most articles (167, 56%) were accessible only through a paywall (Figure 1). Google Scholar searches on public internet connections provided full-text access to 125 articles (42%), and another 83 (28%) were available through institutional access at Oklahoma State University. This highlights that large portions of the surgical literature are unavailable to surgeons or even researchers with broad academic privileges.

**Figure 1.**
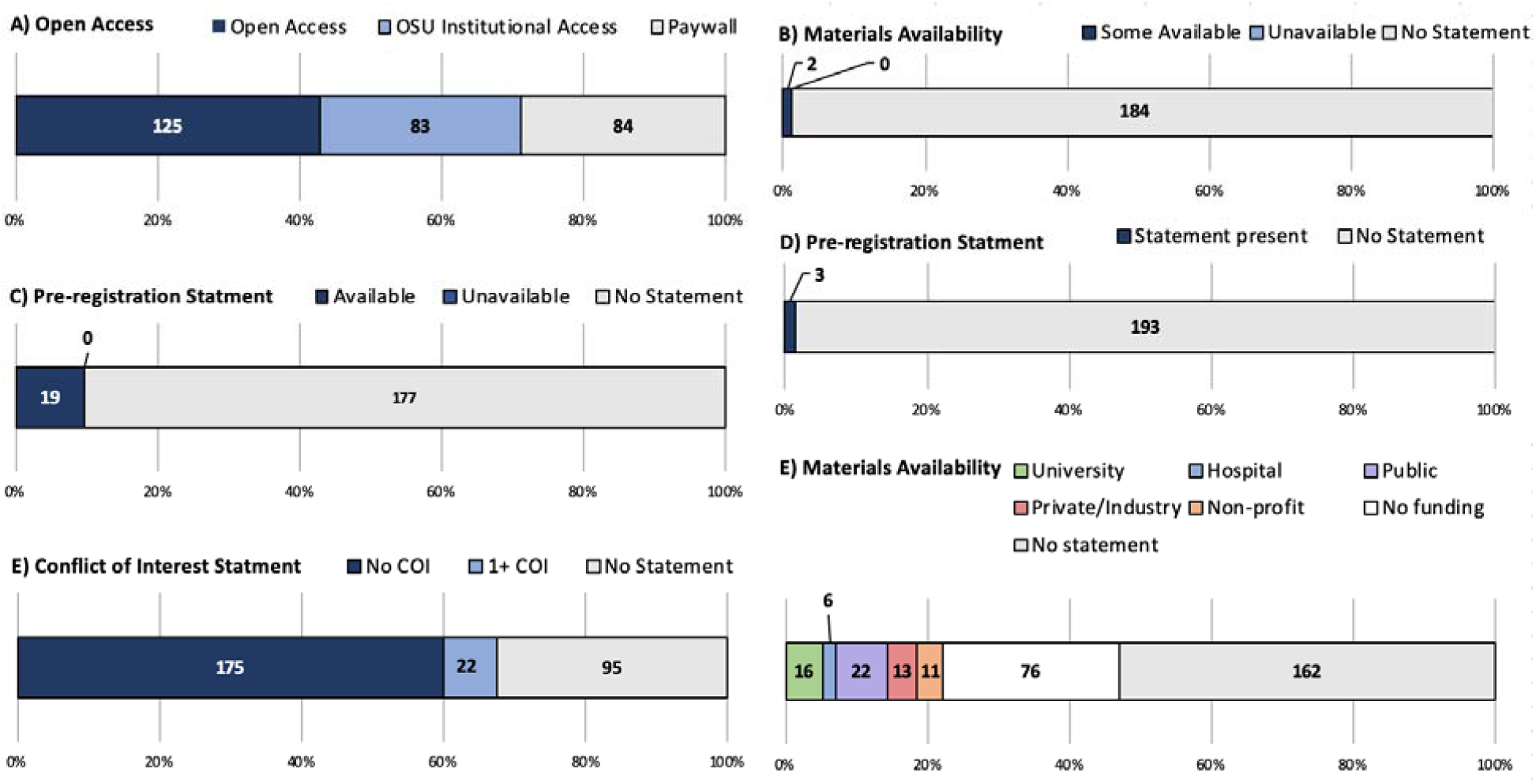

### Availability of Materials and Protocols

Virtually all (184, 99%) articles with empirical data did not indicate any materials necessary to replicate their study. Of the two that provided materials statements, one stated that the product used was commercially available, and the other was a table containing ICD-9 coding information. Only one study provided a protocol, in the form of a registration number, which could not be found on the journal’s website.

### Availability of Data

Nineteen articles (9.7%) indicated data were available, with the distribution of access shown in Figure 2. Most of these could be accessed and contained clearly documented data files, whereas only six (3.1%) contained all the raw data necessary to reproduce the findings reported in the study.

**Figure 2.**
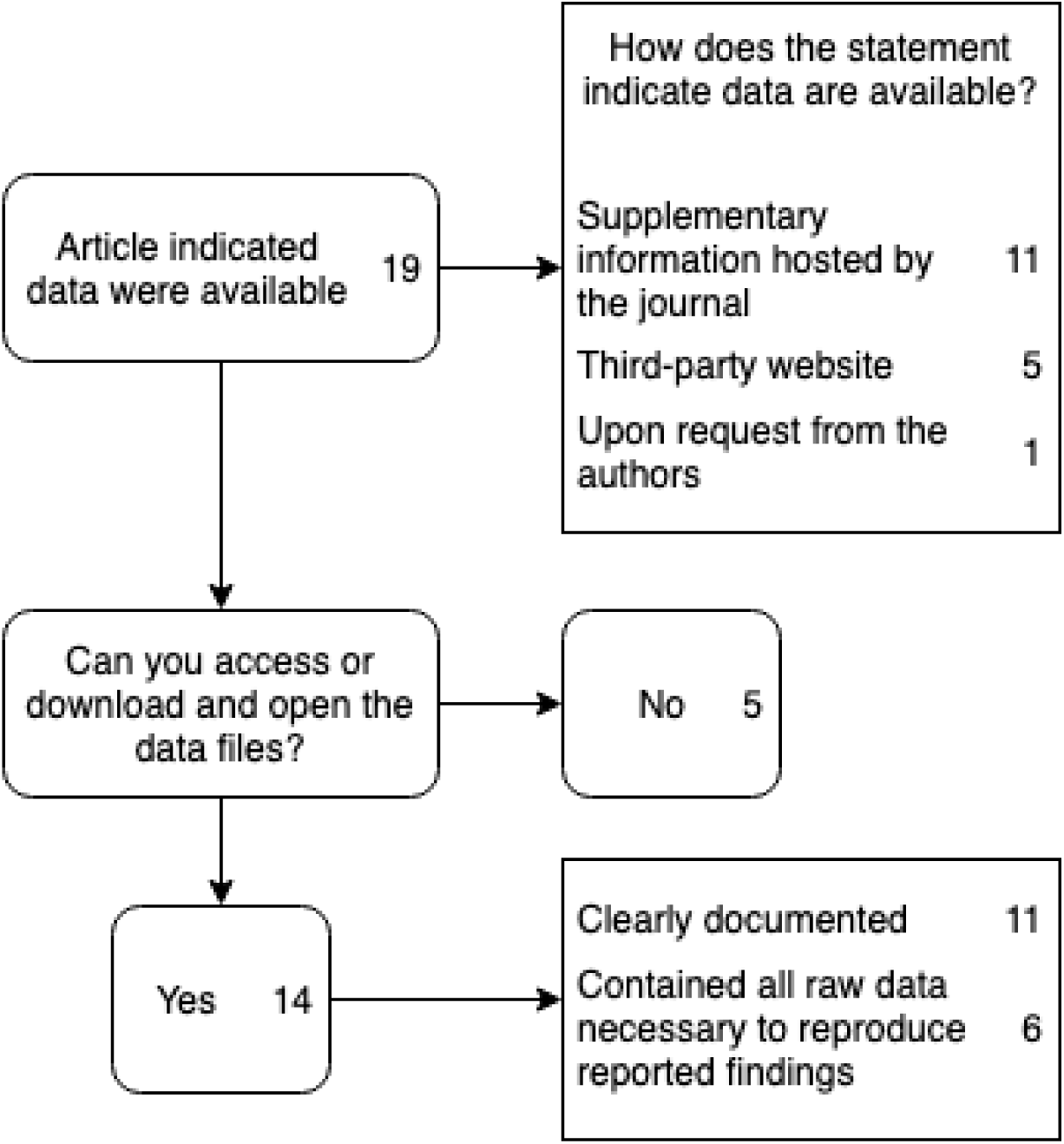

### Availability of Analysis Scripts

Of the 197 articles with empirical data, zero indicated that analysis scripts were available. This demonstrates the limitations of published data in surgical journals, even when raw data were available.

### Availability of Preregistration Information

Preregistration allows researchers to describe their hypotheses, methods, and analyses before research is conducted, in a way that can be externally verified to avoid bias^18^. A large majority of articles (193, 98%) articles did not contain a statement concerning preregistration. Only three provided a related statement, of which two could be accessed and only one contained an explicit hypothesis and methods outline.

### Availability of Conflicts of Interest and Funding Statements

Most (197, 68%) articles provided a statement concerning conflicts of interest. Twenty-two of these (7.5%) declared one or more conflicts of interest, and 95 (33%) did not include a conflict of interest statement. Funding sources were slightly more obscure, with 162 (55.5%) declining to provide a funding statement. Of those citing funding, 52.8% stated that their research was not explicitly funded. Public (e.g., NIH) and university funding were most commonly reported in the surgical literature (Figure 1).

## Discussion

Our assessment of 300 randomly selected surgery publications indicates that the current body of research lacks basic elements for transparent and reproducible practices. One fundamental principle of the scientific method is that experiments must be reproducible. Scientific validity and knowledge increase when researchers can replicate results from previous experimentation. If an experiment or study can be independently verified, its conclusions can be considered more reliable.^2^ Our results strengthen the current consensus in meta-research literature that both observational^10,19,20^ and experimental^21,22^ studies in many surgical fields are generally deficient in transparency and reproducibility, though previous research has not applied our sampling techniques to enable generalizability. In this section, we discuss several findings with particularly poor results.

## Materials and Protocol

The availability of study materials and protocols is fundamental to the reliability of reported outcomes. This reproducibility problem is made worse when funding sources and publishers overvalue high-profile journals and incentivize publication quantity over quality.^23^ The NIH has developed a training program to address this concern,^24^ though the responsibility often falls on journals to require these elements of reproducibility. Steward et al. describes the economic pressures on scientific journals to truncate the methods section of manuscripts or move it to supplementary information, noting that the section rarely contains enough detail for an exact replication.^25^ For example, in surgical literature, this protocol should include a careful description of the surgical plan, intraoperative decision making, case variability, and enough detail to standardize the patients’ pre- and post-operative medical care. Our analysis validated the hypothesis that few surgical manuscripts provide materials or protocols.^26^ These findings highlight a costly inefficiency gap in surgical research. It is fundamentally impossible to build on the foundation of surgical knowledge without enough detail to replicate published results. Clinical research is expensive, and confirming results with high-quality replications is even more costly when the original detail is missing and multiple iterations of a protocol are required to complete the replication.^27,28^

### Open Access

The availability, or open access, of a study directly influences the proportion of health professionals who will read and make decisions based on the study. Previous research has shown that articles in clinical medicine are less likely to have open access, compared to those in other fields of study.^29^ For example, in a study by Hardwicke et al., only 95 of 198 studies performed in three hospital systems were publicly accessible.^30^ For example, in a study by Hardwicke et al., only 95 of 198 studies performed in three hospital systems were publicly accessible. In our study, we found that less than half of the articles were available through our university’s broad institutional access. Thus, smaller universities and unaffiliated researchers who lack comprehensive library subscriptions are at a disadvantage when trying to reproduce a study. Open access should transcend academic affiliation to support sustainable lifelong learning.^31^ Yet, we found that only 42.6% of the articles were available publicly, and 28.6% were not accessible at all. This finding suggests that researchers and physicians lack access to pertinent information relevant to their specialties. Further, paywall restrictions may create preferential disadvantages among rural or hospital-based surgeons who lack affiliations to an academic institution, because their access is limited to abstracts. A study by Marcello et al. demonstrates that clinical decisions in surgery guided by full-text articles are more accurate than those guided by abstracts alone.^32^ Previous research indicates that abstracts may not be as informative due to the high incidence of spin, a form of selective reporting bias where the results in an abstract are inconsistent with the actual findings in the body of the paper.^33–35^ As such, these paywalls limit the self-correction of the scientific process and spread of the latest medical research, which is deleterious to patient care. By limiting a physician’s ability to effectively analyze the literature, poor clinical decisions become more likely. Self-correction is a necessary safeguard in surgical specialties, limiting poor outcomes and loss of life.

### Pre-registration

The subject of pre-registration has garnered increased interest in the medical community.^4^ Study outcomes are frequently interpreted as broad clinical principles rather than solely applicable to the cross-section of conditions for which an intervention has been tested. Pre-registration directly confronts this form of “hypothesizing after the results are known,” or HARKing,^5^ and discourages the use of narrow, context-bound results in broader clinical practice. In our sample, only one randomized controlled trial was adequately pre-registered. Further, of the top-five surgical journals by h-index, none required pre-registration of observational studies.^36–40^ As of 2007, the FDA Amendments Act mandates pre-registration of most clinical trials.^41,42^ A suspiciously sharp fall in positive findings following its implementation suggests a tendency of study authors to engage in selective reporting bias.^43^ Though efforts like the Consolidated Standards of Reporting Trials (CONSORT)^44^ aim at improving the reporting of randomized controlled trials, this tenant of academic transparency does not currently affect most surgical literature, which consists of observational database studies. For example, of the studies in our sample containing empirical data, we found that 93% were observational in design. Further, healthcare database research has been shown to lack complete reporting of study implementation.^23^ Given that pre-registration confronts hindsight bias^4^ and that no gold standard of observational reproducibility exists, it is the opinion of the authors that additional regulation mandating the pre-registration of observational studies would strengthen the field of surgical research.

### Strengths and Weaknesses

Our study has many strengths, including the construction of our random sample, generalizability to other areas of clinical medicine, extensive training of our coders, and use of best practices of meta-research. The blinded, double-extraction technique implemented in this study is considered the gold standard for meta-research data extraction,^45,46^ and it increases the validity of our findings. In addition, the robust training provided to investigators prior to the start of coding adds reliability to the study results. We also note some important limitations. First, we acknowledge the inherent possibility of sampling error, though the authors maintain the generalizability of our findings, as we used a random sampling procedure. Second, no authors were contacted to request additional information, so our outcomes are based on published research alone. According to one study on highly cited articles, 68% of contacted authors either did not respond or would not share the data, 18% said the data was partially available, and only 14% offered unrestricted access to the data.^30^

Lack of reproducibility increases development costs of drugs, medical devices, protocols, and procedures. Most important, it can have deleterious effects on patient health in every stage of surgical research, leading to increased risk of treatment failure.^4^ It is our hope that study authors improve their reporting standards to increase the transparency and reproducibility of surgical research in the name of better science, better outcomes, and progression of the field of surgery.

## Funding Statement

This study has no funding.

## Conflict of Interest Statement

All authors declare no conflicts of interest.

## Supplement

**Supplemental Table 1:** Sample Characteristics of Surgery Literature.

| **Characteristic** |  | **Variables** |  |
| --- | --- | --- | --- |
|  |  | **N (%)** | **95% CI** |
| **Test Subjects**<br>**N=292** | Animals | 7 (2.40) | 0.007-0.041 |
|  | Humans | 200 (68.5) | 0.632-0.737 |
|  | Both | 0 | 0 |
|  | Neither | 85 (29.1) | 0.240-0.343 |
| **Country of Journal Publication**<br>**N=292** | US | 210 (71.9) | 0.668-0.770 |
|  | UK | 44 (15.1) | 0.110-0.191 |
|  | Germany | 6 (2.05) | 0.004-0.037 |
|  | China | 4 (1.37) | 0.001-0.027 |
|  | Italy | 4 (1.37) | 0.001-0.027 |
|  | France | 3 (1.03) | 0.000-0.022 |
|  | Other | 20 (6.85) | 0.040-0.097 |
| **Country of Corresponding Author**<br>**N=292** | US | 112 (38.1) | 0.326-0.436 |
|  | Japan | 18 (6.12) | 0.034-0.088 |
|  | UK | 17 (5.78) | 0.031-0.084 |
|  | China | 16 (5.44) | 0.029-0.080 |
|  | Italy | 12 (4.08) | 0.018-0.063 |
|  | France | 11 (3.74) | 0.016-0.059 |
|  | Canada | 10 (3.40) | 0.014-0.055 |
|  | Spain | 10 (3.40) | 0.014-0.055 |
|  | Germany | 7 (2.38) | 0.007-0.041 |
|  | India | 2 (0.68) | 0.000-0.016 |
|  | Other | 62 (21.68) | 0.170-0.263 |
| **Open Access**<br>**N=292** | Open access | 125 (41.7) | 0.361-0.472 |
|  | OSU<br>institutional<br>access | 83 (27.7) | 0.226-0.327 |
|  | Paywall | 84 (28.0) | 0.229-0.331 |
| **5-Year Impact**<br>**Factor N=292** | Median | 2.727 | - |
|  | 1st quartile | 1.788 | - |
|  | 3rd quartile | 3.798 | - |
|  | Interquartile<br>range | 2.108-3.533 | - |
| **Most Recent**<br>**Impact Factor**<br>**Year N=300** | 2014 | 0 | - |
|  | 2015 | 0 | - |
|  | 2016 | 0 | - |
|  | 2017 | 268 | - |
|  | 2018 | 19 | - |
|  | Not found | 13 | - |
| **Most Recent**<br>**Impact Factor**<br>**N=286** | Median | 2.471 | - |
|  | 1st quartile | 1.586 | - |
|  | 3rd quartile | 3.621 | - |
| Abbreviations: CI, confidence interval |  |  |  |

**Supplemental Table 2:**
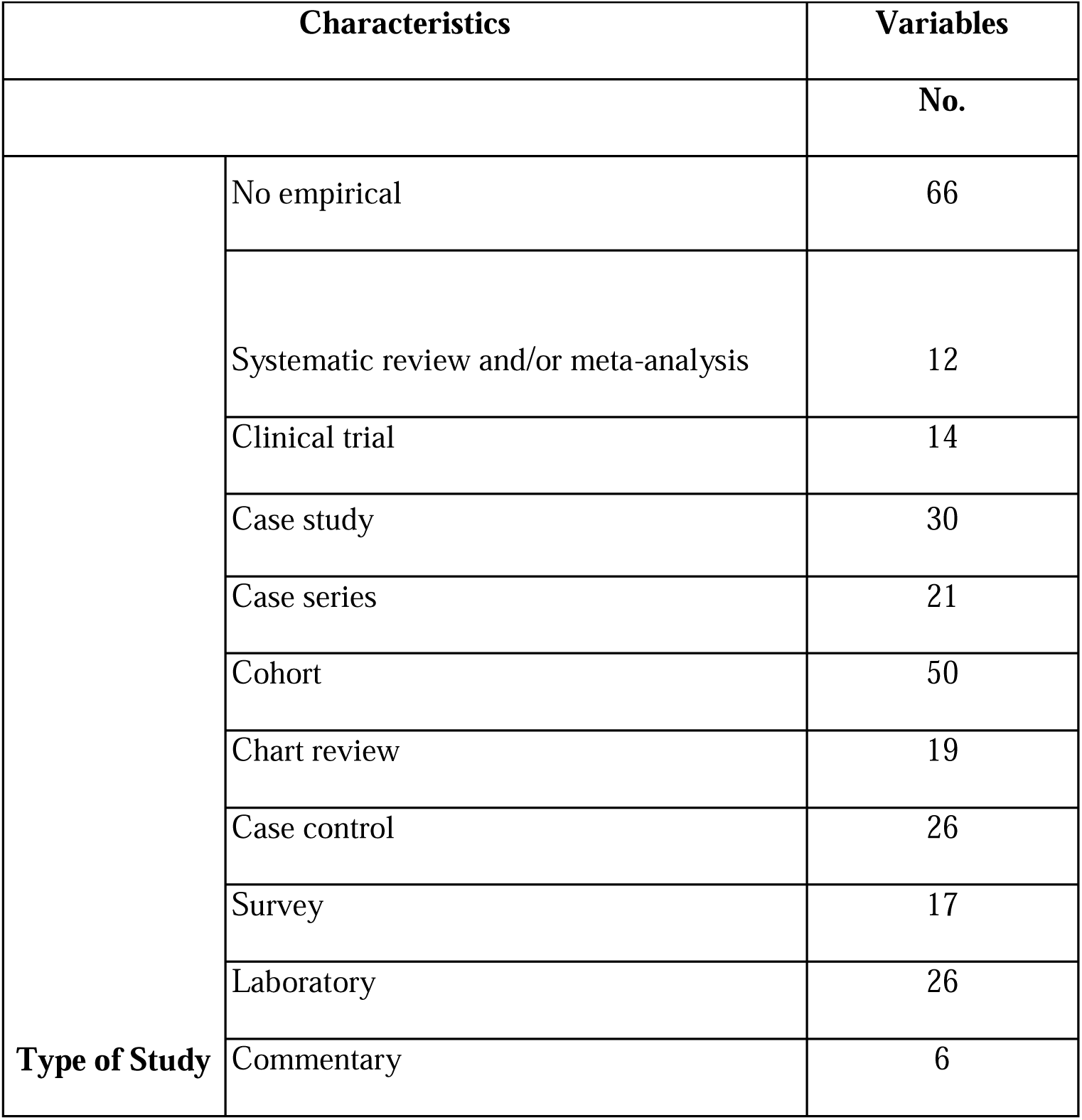
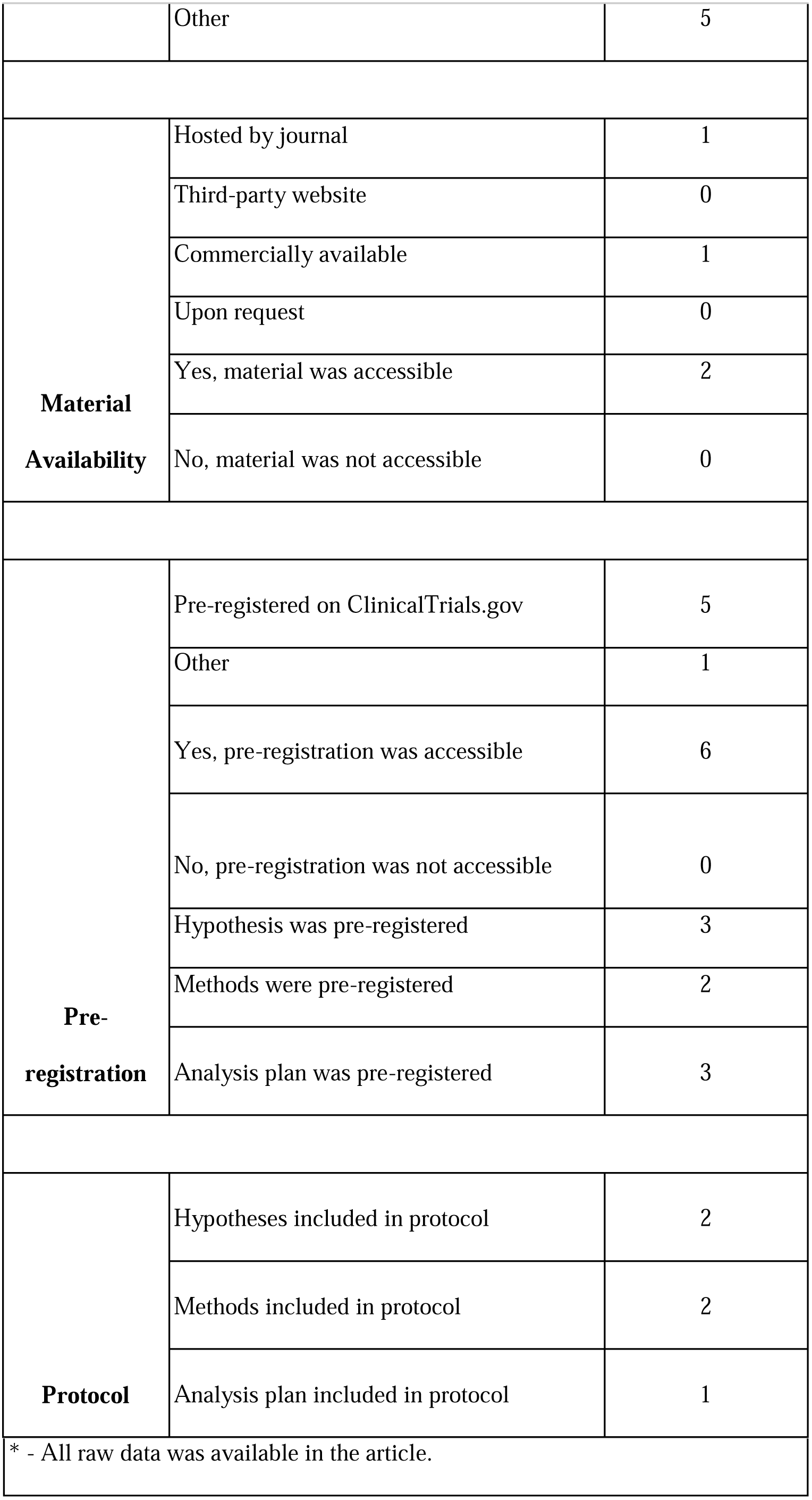
Additional Reproducibility Characteristics.

| **Supplemental Table 2: Additional Reproducibility Characteristics** |  |  |
| --- | --- | --- |
| **Characteristics** |  | **Variables** |
|  |  | **No.** |
| **Type of Study** | No empirical | 66 |
|  | Systematic review and/or meta-analysis | 12 |
|  | Clinical trial | 14 |
|  | Case study | 30 |
|  | Case series | 21 |
|  | Cohort | 50 |
|  | Chart review | 19 |
|  | Case control | 26 |
|  | Survey | 17 |
|  | Laboratory | 26 |
|  | Commentary | 6 |
|  | Other | 5 |
| **Material Availability** | Hosted by journal | 1 |
|  | Third-party website | 0 |
|  | Commercially available | 1 |
|  | Upon request | 0 |
|  | Yes, material was accessible | 2 |
|  | No, material was not accessible | 0 |
| **Pre-registration** | Pre-registered on ClinicalTrials.gov | 5 |
|  | Other | 1 |
|  | Yes, pre-registration was accessible | 6 |
|  | No, pre-registration was not accessible | 0 |
|  | Hypothesis was pre-registered | 3 |
|  | Methods were pre-registered | 2 |
|  | Analysis plan was pre-registered | 3 |
| **Protocol** | Hypotheses included in protocol | 2 |
|  | Methods included in protocol | 2 |
|  | Analysis plan included in protocol | 1 |

\* - All raw data was available in the article.

## References

1. Goodman, S. N., Fanelli, D. & Ioannidis, J. P. A. What does research reproducibility mean? Sci. Transl. Med. 8, 341ps12 (2016).

2. Downing, S. M. Reliability: on the reproducibility of assessment data. Med. Educ. 38, 1006–1012 (2004).

3. Chalmers, I. & Glasziou, P. Avoidable waste in the production and reporting of research evidence. Lancet 374, 86–89 (2009).

4. Nosek, B. A., Ebersole, C. R., DeHaven, A. C. & Mellor, D. T. The preregistration revolution. Proc. Natl. Acad. Sci. U. S. A. 115, 2600–2606 (2018).

5. Kerr, N. L. HARKing: hypothesizing after the results are known. Pers. Soc. Psychol. Rev. 2, 196–217 (1998).

6. Baker, M. 1,500 scientists lift the lid on reproducibility. Nature 533, 452–454 (2016).

7. Plan to replicate 50 high-impact cancer papers shrinks to just 18. Science | AAAS (2018). Available at: https://www.sciencemag.org/news/2018/07/plan-replicate-50-high-impact-cancer-papers-shrinks-just-18. (Accessed: 20th July 2019)

8. Reproducibility Project: Cancer Biology. Available at: https://cos.io/rpcb/. (Accessed: 20th July 2019)

9. Open Science Collaboration. PSYCHOLOGY. Estimating the reproducibility of psychological science. Science 349, aac4716 (2015).

10. Agha, R. A. et al. The STROCSS statement: Strengthening the Reporting of Cohort Studies in Surgery. Int. J. Surg. 46, 198–202 (2017).

11. Agha, R. A. et al. Preferred reporting of case series in surgery; the PROCESS guidelines. Int. J. Surg. 36, 319–323 (2016).

12. Framework - The IDEAL Collaboration. Available at: http://www.ideal-collaboration.net/framework/. (Accessed: 20th July 2019)

13. Lawton, J. S. Reproducibility and replicability of science and thoracic surgery. J. Thorac. Cardiovasc. Surg. 152, 1489–1491 (2016).

14. Ascha, M., Ascha, M. S. & Gatherwright, J. The Importance of Reproducibility in Plastic Surgery Research. Plast. Reconstr. Surg. 144, 242–248 (2019).

15. Hardwicke, T. E., Wallach, J. D., Kidwell, M. & Ioannidis, J. An empirical assessment of transparency and reproducibility-related research practices in the social sciences (2014-2017). (2019). doi:10.31222/osf.io/6uhg5

16. Murad, M. H. & Wang, Z. Guidelines for reporting meta-epidemiological methodology research. Evid. Based. Med. 22, 139–142 (2017).

17. of Health, U. S. D., Services, H. & Others. CFR 46. Public welfare protection of human subjects. http://www.hhs.gov/ohrp/humansubjects/guidance/45cfr46.htm 46, (45AD).

18. van’t Veer, A. E. & Giner-Sorolla, R. Pre-registration in social psychology—A discussion and suggested template. J. Exp. Soc. Psychol. 67, 2–12 (2016).

19. Agha, R. A., Lee, S.-Y., Jeong, K. J. L., Fowler, A. J. & Orgill, D. P. Reporting Quality of Observational Studies in Plastic Surgery Needs Improvement: A Systematic Review. Ann. Plast. Surg. 76, 585–589 (2016).

20. Sorensen, A. A., Wojahn, R. D., Manske, M. C. & Calfee, R. P. Using the Strengthening the Reporting of Observational Studies in Epidemiology (STROBE) Statement to assess reporting of observational trials in hand surgery. J. Hand Surg. Am. 38, 1584–9.e2 (2013).

21. Agha, R. A. et al. Randomised controlled trials in plastic surgery: a systematic review of reporting quality. Eur. J. Plast. Surg. 37, 55–62 (2014).

22. Ghert, M. The reporting of outcomes in randomised controlled trials: The switch and the spin. Bone Joint Res. 6, 600–601 (2017).

23. Franzoni, C., Scellato, G. & Stephan, P. Science policy. Changing incentives to publish. Science 333, 702–703 (2011).

24. Collins, F. S. & Tabak, L. A. Policy: NIH plans to enhance reproducibility. Nature 505, 612–613 (2014).

25. Steward, O., Popovich, P. G., Dietrich, W. D. & Kleitman, N. Replication and reproducibility in spinal cord injury research. Exp. Neurol. 233, 597–605 (2012).

26. Pidgeon, T. E. et al. The use of study registration and protocols in plastic surgery research: A systematic review. Int. J. Surg. 44, 215–222 (2017).

27. Coles, N., Tiokhin, L., Scheel, A. M., Isager, P. M. & Lakens, D. The Costs and Benefits of Replication Studies. (2018). doi:10.17605/OSF.IO/C8AKJ

28. Freedman, L. P., Cockburn, I. M. & Simcoe, T. S. The Economics of Reproducibility in Preclinical Research. PLoS Biol. 13, e1002165 (2015).

29. Iqbal, S. A., Wallach, J. D., Khoury, M. J., Schully, S. D. & Ioannidis, J. P. A. Reproducible Research Practices and Transparency across the Biomedical Literature. PLoS Biol. 14, e1002333 (2016).

30. Hardwicke, T. E. & Ioannidis, J. P. A. Populating the Data Ark: An attempt to retrieve, preserve, and liberate data from the most highly-cited psychology and psychiatry articles. PLoS One 13, e0201856 (2018).

31. Tennant, J. P. et al. The academic, economic and societal impacts of Open Access: an evidence-based review. F1000Res. 5, 632 (2016).

32. Marcelo, A. et al. A comparison of the accuracy of clinical decisions based on full-text articles and on journal abstracts alone: a study among residents in a tertiary care hospital. Evid. Based. Med. 18, 48–53 (2013).

33. Austin, J. et al. Evaluation of spin within abstracts in obesity randomized clinical trials: A cross-sectional review. Clin. Obes. 9, e12292 (2019).

34. Jellison, S. et al. Evaluation of spin in abstracts of papers in psychiatry and psychology journals. BMJ Evid Based Med (2019). doi:10.1136/bmjebm-2019-111176

35. Cooper, C. M. et al. Evaluation of spin in the abstracts of otolaryngology randomized controlled trials. Laryngoscope (2018). doi:10.1002/lary.27750

36. Editorial Manager - Annals of Surgery. Available at: http://edmgr.ovid.com/annsurg/accounts/ifauth.htm. (Accessed: 16th August 2019)

37. Information for Authors. Available at: https://www.editorialmanager.com/jvs/account/Information%20for%20authors.html. (Accessed: 16th August 2019)

38. Instructions for Authors | JAMA Surgery | JAMA Network. Available at: https://jamanetwork.com/journals/jamasurgery/pages/instructions-for-authors. (Accessed: 16th August 2019)

39. Author Guidelines - BJS. Available at: https://onlinelibrary.wiley.com/page/journal/13652168/homepage/forauthors.html.

40. Instructions for Authors. (2019). Available at: http://www.springer.com/cda/content/document/cda_downloaddocument/Instructions_June2019_web_links.pdf?SGWID=0-0-45-1657716-p29577513.

41. Office of the Commissioner. Food and Drug Administration Amendments Act (FDAAA) of 2007. U.S. Food and Drug Administration (2018). Available at: http://www.fda.gov/regulatory-information/selected-amendments-fdc-act/food-and-drug-administration-amendments-act-fdaaa-2007. (Accessed: 14th August 2019)

42. FDAAA 801 and the Final Rule - ClinicalTrials.gov. Available at: https://clinicaltrials.gov/ct2/manage-recs/fdaaa. (Accessed: 14th August 2019)

43. Woolston, C. Registered clinical trials make positive findings vanish. Nature News 524, 269 (2015).

44. Altman, D. G. et al. The revised CONSORT statement for reporting randomized trials: explanation and elaboration. Ann. Intern. Med. 134, 663–694 (2001).

45. da Costa, B. R. & Juni, P. Systematic reviews and meta-analyses of randomized trials: principles and pitfalls. Eur. Heart J. 35, 3336–3345 (2014).

46. Higgins, J. P. T. & Green, S. Cochrane Handbook for Systematic Reviews of Interventions. (John Wiley & Sons, 2011).

